# Transplantation Elicits a Clonally Diverse CD8^+^ T Cell Response that is Comprised of Potent CD43^+^ Effectors

**DOI:** 10.1101/2022.05.21.492934

**Authors:** Gregory A. Cohen, Melissa A. Kallarakal, Sahana Jayaraman, Francis I. Ibukun, Katherine P. Tong, Linda D. Orzolek, H. Benjamin Larman, Scott M. Krummey

## Abstract

**SUMMARY:** CD8^+^ T cells mediate acute rejection of allografts, which threatens the long-term survival of transplanted organs. The factors that govern differentiation of graft-directed effector CD8^+^ T cells could lead to targeted approaches to limit acute rejection. Using MHC Class I tetramers, we found that allogeneic CD8^+^ T cells were present at an elevated precursor frequency in naïve mice, only modestly increased in number after grafting, and maintained high T cell receptor diversity throughout the immune response. While antigen-specific effector CD8^+^ T cells poorly express the canonical effector marker KLRG-1, expression of the activated glycoform of CD43 defined potent effectors after transplantation. Activated CD43^+^ effector T cells maintained high expression of ICOS in the presence of CTLA-4 Ig, and dual CTLA-4 Ig/anti-ICOS treatment prolonged graft survival. These data demonstrate that graft-specific CD8^+^ T cells have a distinct response profile relative to anti-pathogen CD8^+^ T cells, and that CD43 and ICOS are critical surface receptors that define potent effector CD8^+^ T cell populations that form after transplantation.

## INTRODUCTION

Activation of CD8^+^ T cells relies on the recognition of cognate antigen in the presence of both coreceptor signaling and the local cytokine milieu (Chung et al., 2021; Ford, 2016; Jameson and Masopust, 2018). The balance of these factors shapes the magnitude and character of the CD8^+^ T cell response. In transplantation, CD8^+^ T cells can respond to allogeneic antigen presented on donor MHC Class I proteins and mediate tissue damage (Callemeyn et al., 2021; Felix et al., 2007). Recent clinical studies have highlighted that the occurrence of acute rejection has cumulative and detrimental effects on the long-term survival of transplanted organs (Cherukuri et al., 2021; Cole et al., 2008; Meier-Kriesche et al., 2000).

Activated T cells undergo several well-established phenotypic changes that distinguish them from quiescent naïve CD8^+^ T cells. The majority of the work underlying these paradigms has been performed in infection models, which present distinct profiles of inflammation and antigen relative to transplanted organs (Braza et al., 2016; Colvin et al., 2017; Dwyer and Turnquist, 2021). The canonical phenotype of an effector CD8^+^ T cell, CD44^hi^CD62L^lo^, does not specifically distinguish the actively responding graft-specific CD8^+^ T cells among the larger pool of previously activated cells. Expression of the surface receptor KLRG-1 is also used to identify actively responding effector CD8^+^ T cells relative to memory precursor cells (Herndler-Brandstetter et al., 2018a; Joshi et al., 2007; Sarkar et al., 2008). However, KLRG-1 expression is not functionally required for the effector function of CD8^+^ T cells nor is it induced after all types of priming *in vivo* (Bozeman et al., 2018; Joshi et al., 2007). Thus, in order to more granularly understand anti-graft CD8^+^ T cell responses and develop strategies to target the allogeneic response, a deeper understanding of the differentiation program of post-graft CD8^+^ T cells is needed.

Due to the technical difficulties associated with identifying allogeneic peptide epitopes, the CD8^+^ T cell response is typically evaluated by using either TCR transgenic mice (e.g. OT-I/OVA) or evaluating the “bulk” pool of activated CD8^+^ T cells. Both approaches sacrifice potential insights gained by assessing the acutely responding antigen-specific T cell clones in the context of physiologic antigen levels, precursor frequency, clonal competition, and inflammatory signals. MHC tetramers have provided a tremendous advance in the ability to identify and track antigen-specific CD8^+^ T cell responses to pathogens and model antigens (Haluszczak et al., 2009; Jenkins and Moon, 2012; Kotturi et al., 2008; Obar et al., 2008). However, the use of MHC tetramers to study allogeneic T cells has been limited due to the technical challenges of identifying appropriate epitopes from across the proteome.

We sought to develop the use of MHC tetramers in a fully allogeneic transplant model by taking advantage of elegant work performed in the characterization of the allogeneic 2C TCR transgenic mouse model several decades ago (Chen et al., 2003; Sykulev et al., 1994a; Udaka et al., 1992). We found that MHC tetramers of an H-2L^d^ restricted epitope for a ubiquitously expressed metabolic enzyme defined a population of CD8^+^ T cells in H-2^b^ C57Bl/6 mice. This approach allowed us to directly study the CD8^+^ T cell response to fully allogeneic H-2^d^ grafts and characterize the differentiation of effector populations in the context of physiologic antigen and inflammation. Here we report the characteristics of the CD8^+^ T cell response to allogeneic antigen, including the clonal features of antigen-specific alloreactive CD8^+^ T cells, and the emergence of a potent population of effector CD8^+^ T cells that express the surface receptor CD43 after grafting.

## RESULTS

### MHC Class I tetramer can be used to identify L^d^ QL9^+^ CD8^+^ T cells in naïve C57Bl/6 mice

Relatively few allogeneic epitopes have been identified in the fully allogeneic Balb/c (H-2^d^) to C57Bl/6 (H-2^b^) transplant model. The alloreactive 2C TCR recognizes H-2L^d^ restricted peptides for the self-peptide α-ketoglutarate dehydrogenase(Chen et al., 2003; Sykulev et al., 1994a; Udaka et al., 1992), thus representing a directly presented allogeneic epitope. We hypothesized that MHC tetramers of this H-2L^d^ restricted epitope could be used to identify endogenous alloreactive CD8^+^ T cells. In order to reliably detect rare CD8^+^ T cells, we incubated single cell suspensions of secondary lymphoid organs with L^d^ QL9 tetramers and enriched with magnetic beads specific for the fluorophore(s) of the tetramer **(Figure 1A)**(Moon et al., 2007). This approach results in a tetramer enriched (column “bound”) and bulk CD8^+^ (column “unbound”) fractions of cells for analysis.

**Figure 1.**
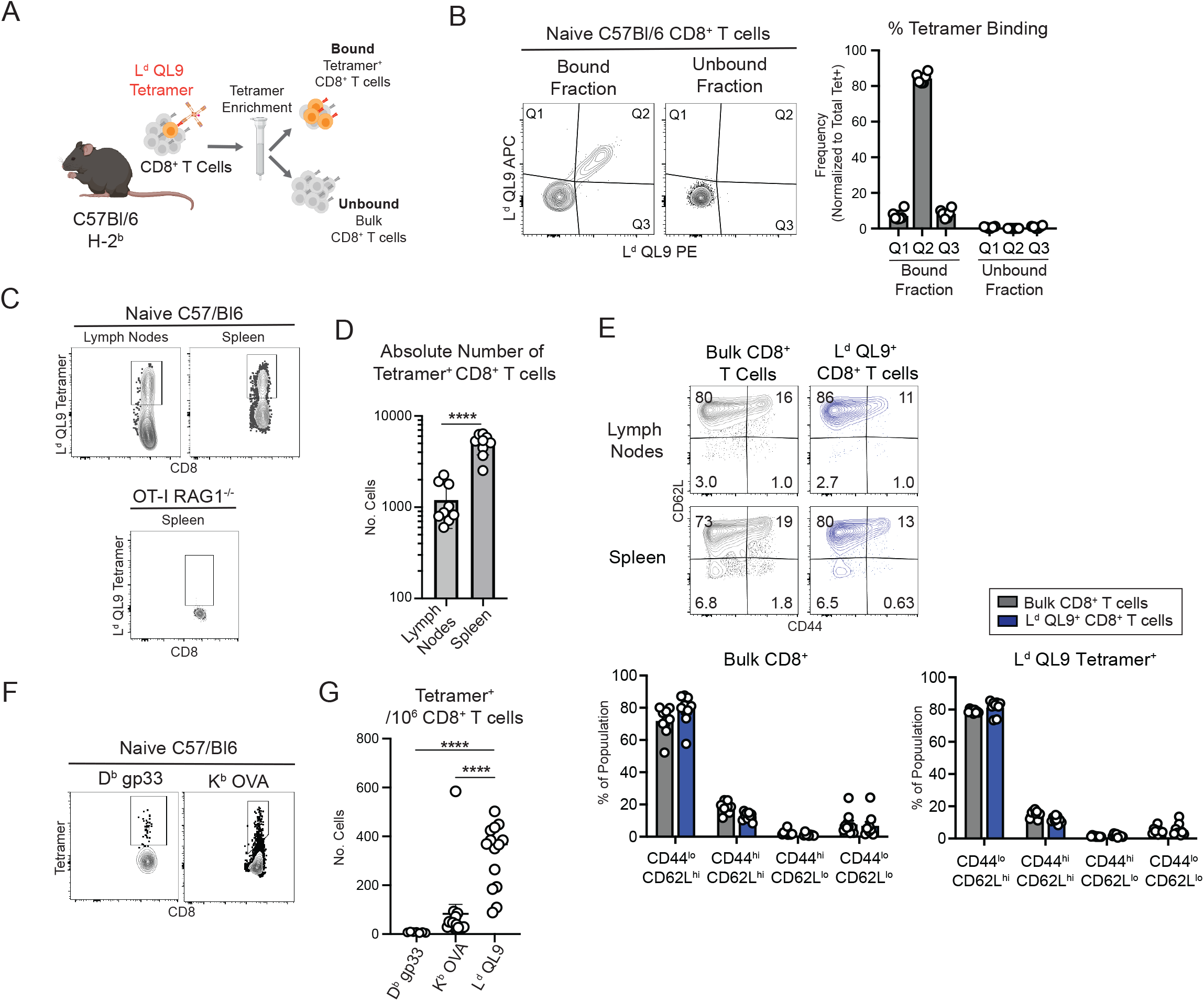
H-2L^d^ QL9 tetramers identify a population of naive CD8^+^ T cells at elevated precursor frequency. (A) Single cell suspensions of lymph node or spleen tissue were incubated with MHC Class I tetramers, and enriched using magnetic beads specific for the tetramer fluorophores, creating column bound tetramer-enriched and unbound bulk CD8^+^ T cell fractions for analysis. (B) Representative flow cytometry plots and summary data depicting the frequency of CD8^+^ T cells binding to L^d^ QL9 PE and L^d^ QL9 APC tetramers in the bound and unbound fractions. (C) Representative flow cytometry plots of L^d^ QL9 tetramer staining in naive C57Bl/6 spleen, C57Bl/6 lymph nodes, or OT-I RAG1^-/-^ spleen. (D) Absolute numbers of L^d^ QL9 tetramer binding CD8^+^ T cells in the lymph nodes and spleen of naive C57Bl/6 mice. (E) Representative flow cytometry plots and summary data depicting the phenotype of L^d^ QL9 tetramer binding CD8^+^ T cells in the lymph nodes and spleen. (F) Representative flow cytometry plots of D^b^ LCMV gp33 and K^b^ OVA tetramer binding CD8^+^ T cells. (G) Number of D^b^ LCMV gp33, K^b^ OVA, and L^d^ QL9 tetramer binding cells per million CD8^+^ T cells. Each point depicts an individual mouse. Summary data depicts pooled results from (B) 3 independent experiments (n=6/group), (D-E) 2 independent experiments (n=9/group) and (G) 2 independent experiments (n=7-14/group). Statistical analyses performed using (D) Student’s unpaired t-test (2-tailed), (G) 1-way ANOVA with Dunnett’s multiple comparisons test. Error bars depict SEM. ****p<0.0001.

In naïve C57Bl/6 mice, we detected a population of L^d^ QL9 tetramer binding cells using PE and APC tetramers **(Figure 1B)**. Using this approach, we identified L^d^ QL9-specific CD8^+^ T cells in the spleen and lymph nodes (mean 4,994 and 1,199 cells/tissue, respectively**; Figure 1C-D)**. Tetramer binding was specific, as no L^d^ QL9 binding cells were found in OT-I RAG1 KO mice **(Figure 1C)**. The phenotype of L^d^ QL9 binding CD8^+^ T cells was largely CD44^lo^CD62L^hi^, similar to the bulk CD8^+^ T cell population in naive **(Figure 1E)**. We found that the precursor frequency of K^b^ OVA and D^b^ LCMV gp33-specific CD8^+^ T cells, while consistent with prior publications (Jenkins and Moon, 2012), were significantly lower than that of L^d^ QL9 CD8^+^ T cells **(Figure 1F-G)**. Thus, the L^d^ QL9 tetramer specifically binds to a population of naïve CD8^+^ T cells that appear to be at a relatively elevated precursor frequency compared with other T cell populations.

### L^d^ QL9^+^ CD8^+^ T cells become activated and proliferate after H-2^d^ Balb/c skin grafts

Having established that a population of H-2^b^ C57Bl/6 CD8^+^ T cells recognize L^d^ QL9 tetramers in naïve mice, we next wanted to evaluate whether L^d^ QL9^+^ CD8^+^ T cells respond when presented with allogeneic H-2L^d^ antigen-expressing tissue. We grafted skin from H-2^d^ Balb/c mice onto C57Bl/6 mice and assessed tetramer binding CD8^+^ T cells **(Figure 2A)**. We found that a significant portion of L^d^ QL9^+^ CD8^+^ T cells became CD44^hi^ on day 10 after skin grafting in the draining lymph nodes and spleen **(Figure 2B)**, indicative of priming after exposure to cognate antigen. The absolute number of CD44^hi^ L^d^ QL9 tetramer binding cells increased on day 10 in both the draining lymph nodes and spleen **(Figure 2C)**, representing an average 10-fold increase in the draining lymph node and 6.1-fold in the spleen **(Figure 2D)**.

**Figure 2.**
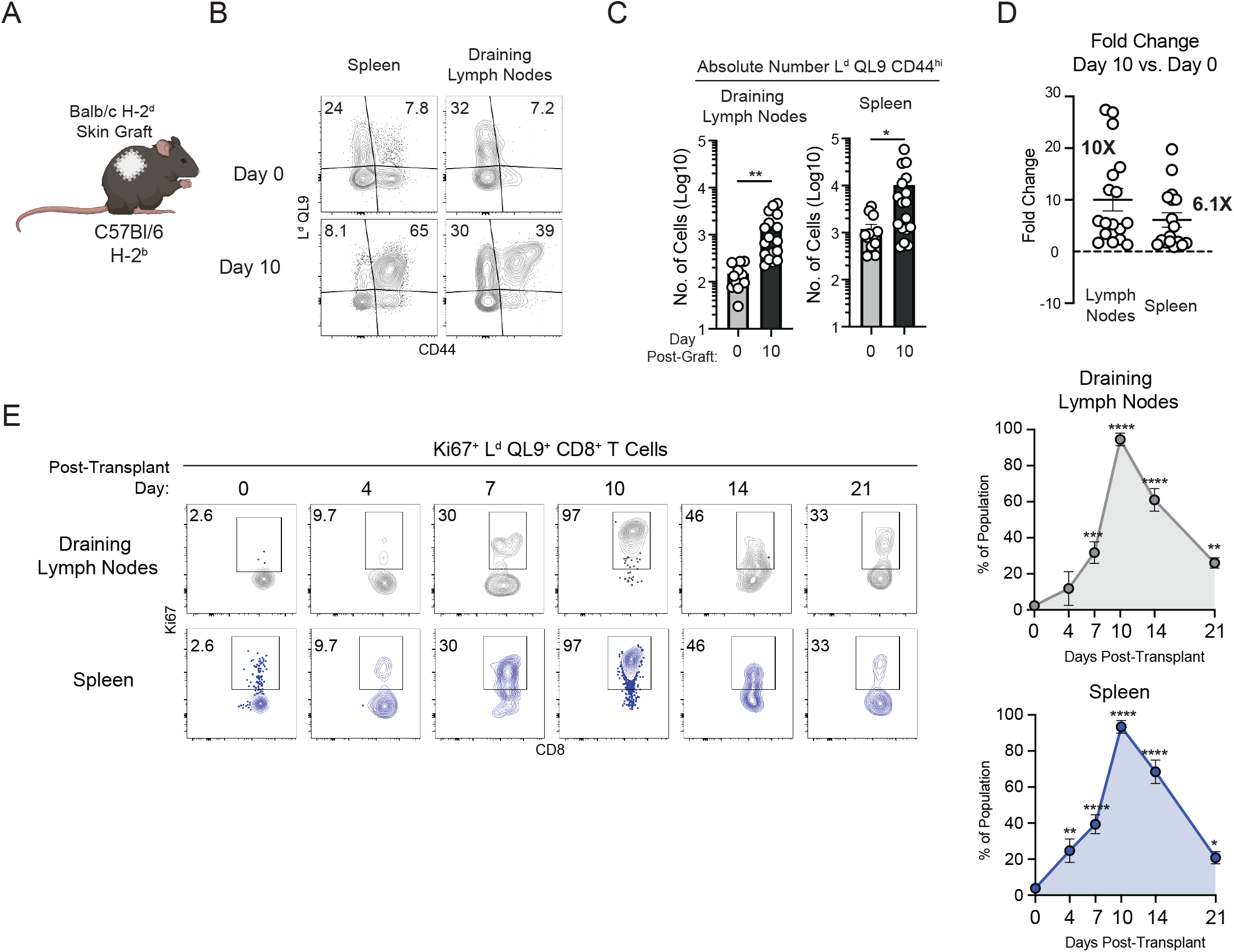
H-L^d^ QL9^+^ CD8^+^ T cells proliferate in the spleen and draining lymph nodes after grafting. (A) C57Bl/6 mice were grafted with H-2^d^ Balb/c skin. (B) Representative flow cytometry plots and (C) absolute numbers of CD8^+^ T cells on day 0 or day 10 after grafting in the draining lymph nodes and spleen. (D) Fold change of the number of L^d^ QL9^+^ CD8^+^ T cells on day 10 post-graft relative to day 0 in the draining lymph nodes or spleen. (E) Representative flow cytometry plots and summary data of Ki67 staining among CD44^hi^ L^d^ QL9^+^ CD8^+^ T cell populations in the draining lymph nodes and spleen. Each data point depicts an individual mouse. Summary data depicts pooled data from (C-D) 4 independent experiments (n=12-17/group) or (E) 2-3 independent experiments (6-9/group). Statistical analyses performed using (C) unpaired Student’s t-test (2-tailed), (D) one-way ANOVA with Dunnett’s multiple comparisons test. Error bars depict SEM. *p<0.05, **p<0.01, ***p<0.001, ****p<0.0001.

In order to assess whether the phenotypic and cell number changes reflected a proliferative response in L^d^ QL9-specific CD8^+^ T cells, we stained cells for Ki67 in order to assess recent and active proliferation after Balb/c skin grafting. We found that in both the draining lymph node and spleen, the frequency of Ki67^+^ cells increased by day 7, and peaked on day 10 relative to naïve CD8^+^ T cells **(Figure 2E)**. At day 10 post-transplant, nearly all of the L^d^ QL9^+^ CD44^hi^ CD8^+^ T cells were Ki67^+^ in both the draining lymph nodes and spleen **(Figure 2E)**. The frequency of Ki67^+^ CD8^+^ T cells also peaked in the bulk CD8^+^ T cell population at day 10 **(Figure S1)**. Thus, as measured by both absolute numbers and Ki67^+^ frequency, L^d^ QL9 tetramer binding CD8^+^ T cells respond to endogenously presented antigen in Balb/c skin grafts.

### Graft-specific CD8^+^ T cells maintain clonal diversity during the peak effector response

Given that the L^d^ QL9-specific CD8^+^ T cells are found at an elevated precursor frequency relative to CD8^+^ T cell populations specific for non-transplant antigens **(Figure 1G)**, yet underwent a relatively modest fold expansion after grafting **(Figure 2D)**, we sought to evaluate the clonal dynamics of this antigen-specific population. We used a multiplexed *TCRB* sequencing platform in order to evaluate the anti-graft CD8^+^ T cell response in naïve and effector graft-specific CD8^+^ T cells. To identify a graft-specific *TCRB* signature, we sequenced FACS-sorted naïve CD44^lo^ L^d^ QL9^+^ T cells from naive mice and effector CD44^hi^ L^d^ QL9^+^ T cells from mice on day 10 post-Balb/c skin graft to compare with bulk (L^d^ QL9^-^) CD8^+^ T cells sorted into CD44^lo^ naïve and CD44^hi^ effector populations **(Figure 3A)**.

**Figure 3.**
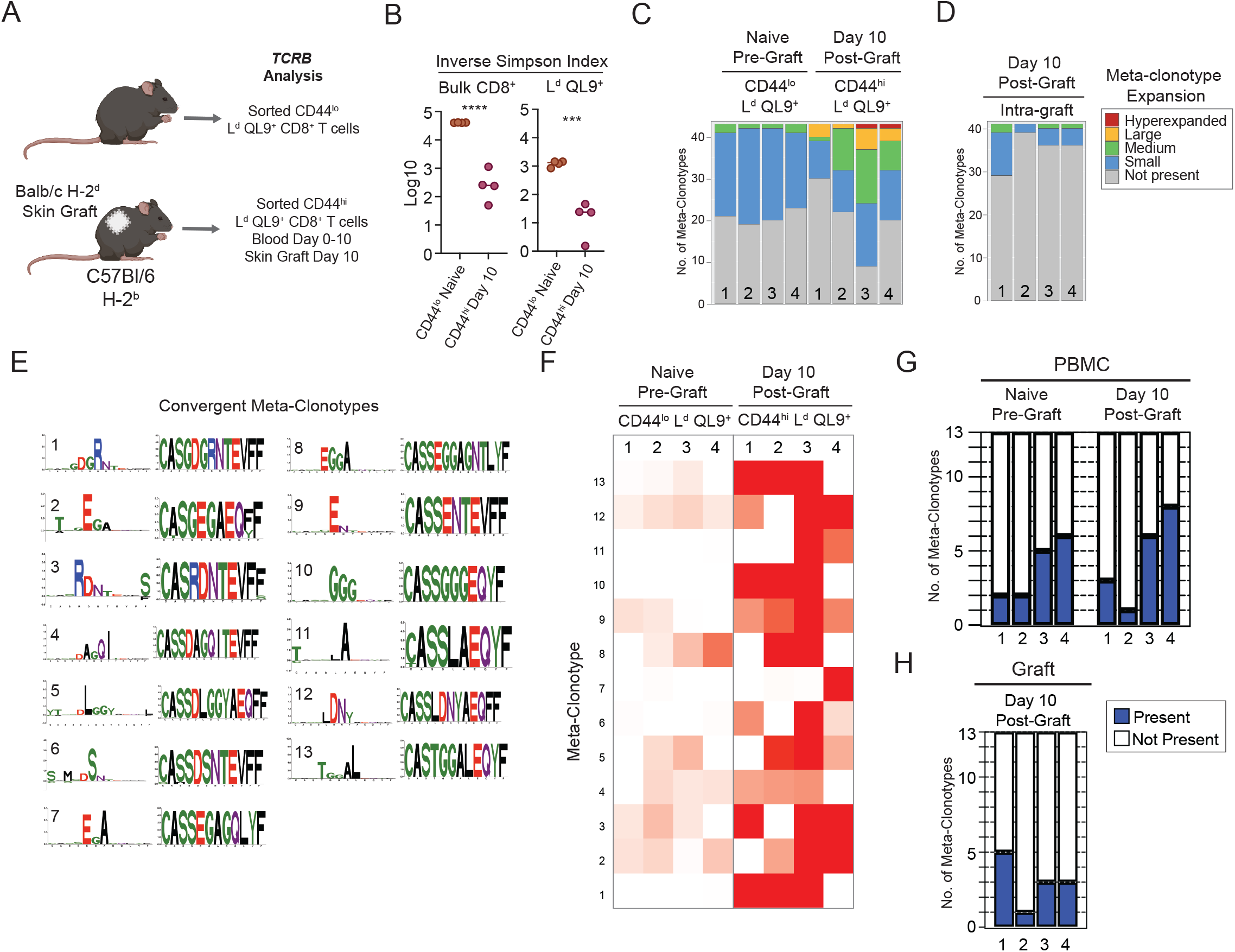
Graft-specific CD8^+^ T cells maintain clonal diversity throughout the primary effector response. (A) C57Bl/6 mice were grafted with Balb/c skin grafts and multiple tissues were collected. Splenic CD8^+^ T cell populations were flow cytometrically sorted based on L^d^ QL9 binding and CD44 expression. (B) Inverse Simpson Index for naïve CD44^lo^ and day 10 effector CD44^hi^ T cell populations are depicted. (C) Frequency of meta-clonotypes in each sample that are defined as hyperexpanded (>1%), large (0.1-1%), medium (0.01-0.1%), small (<0.01%), or are not present. (D) Frequency of meta-clonotypes defined as in (C), found in the skin graft tissue on day 10. (D) CDR3 sequences from 13 convergent meta-clonotypes. Left TCR logo is scaled by per-column relative entropy to background population. The right TCR logo depicts the full meta-clonotype CDR3 sequence. (E) Frequency of meta-clonotypes present in the splenic CD44^lo^ and CD44^hi^ L^d^ QL9^+^ populations. (F-G) Number of meta-clonotypes present in peripheral blood mononuclear cells (PBMC) and graft tissue at the depicted timepoints. Each data point depicts an individual mouse. Statistical analysis performed using unpaired Student’s t-test. Error bars depict SEM. ***p<0.001, ****p<0.0001.

We found that CD44^hi^ L^d^ QL9^+^ T cells had significantly less population diversity versus naïve CD44^lo^ L^d^ QL9^+^ T cells, consistent with greater oligoclonality after grafting **(Figure 3B)**. In order to evaluate the dynamics of clones that were enriched in the CD44^hi^ L^d^ QL9^+^ population, we used the TCRdist package, which identifies biochemically similar CDR3 sequences called meta-clonotypes, in antigen-specific T cell populations evaluated in comparison with background populations (Dash et al., 2017). We identified 43 meta-clonotypes that were enriched over background in our samples **(Table S1)**. Compared with the naïve L^d^ QL9^+^ population, we found that a greater number of the post-graft CD44^hi^ L^d^ QL9^+^ meta-clonotypes were found to have medium expansion, large expansion, and hyper-expansion frequencies **(Figure 3C)**. Interestingly, clones recovered from within the skin grafts on day 10 remained relatively polyclonal, and all grafts contained meta-clonotypes **(Figure 3D)**.

We next evaluated the representation of meta-clonotypes across individual mice, and found that 13 meta-clonotypes were convergent, which we defined as present in 75% or more of the individual mice CD44^hi^ L^d^ QL9^+^ populations **(Figure 3E)**. The defining amino acid sequences of these meta-clonotypes varied, although there was a preference for a glycine at position 5 and/or 6, which was present in 8 of 13 of the convergent meta-clonotypes. We evaluated the expression of the convergent meta-clonotypes and found that they were enriched in the L^d^ QL9^+^ CD8^+^ T cells at day 10 post-graft relative to naïve populations **(Figure 3F)**. In each mouse, at least one of the convergent meta-clonotypes was present **(Figure 3G-H)**. However, we did not find an increased number of convergent meta-clonotypes in the unsorted peripheral blood or skin grafts **(Figure 3G-H)**. Thus, these data demonstrate that after transplantation, CD8^+^ T cells responding to a single allogeneic antigen maintain high clonal diversity. However, there may be biochemical properties that govern CD8^+^ T cell recognition of allogeneic antigen.

### A low frequency of CD8^+^ T_EFF_ express KLRG-1^+^ after grafting

A significant goal in transplantation is to identify acutely responding T cells in order to monitor the risk for, or presence of, acute cellular graft rejection. In order to understand the function of graft-specific effector CD8^+^ T cells, we evaluated the phenotypic profile of L^d^ QL9^+^ effector CD8^+^ T cells (T_EFF_), as defined by CD44^hi^ status. We first evaluated the expression of CXCR3 and CD62L, two surface receptors that have been shown to be important for post-infection T_EFF_ function. We found that nearly all L^d^ QL9^+^ and bulk CD8^+^ T cells became CXCR3^+^ at day 10 and remained stable to a memory timepoint **(Figure 4A-C)**. About half of these CXCR3^+^ T_EFF_ lost CD62L expression, and this frequency of CD62L^lo^ cells was also stable from day 10-42 after skin grafting. Thus, while these markers are useful for identifying antigen-experienced CD8^+^ T cells after transplantation, they do not delineate acutely responding effectors.

**Figure 4.**
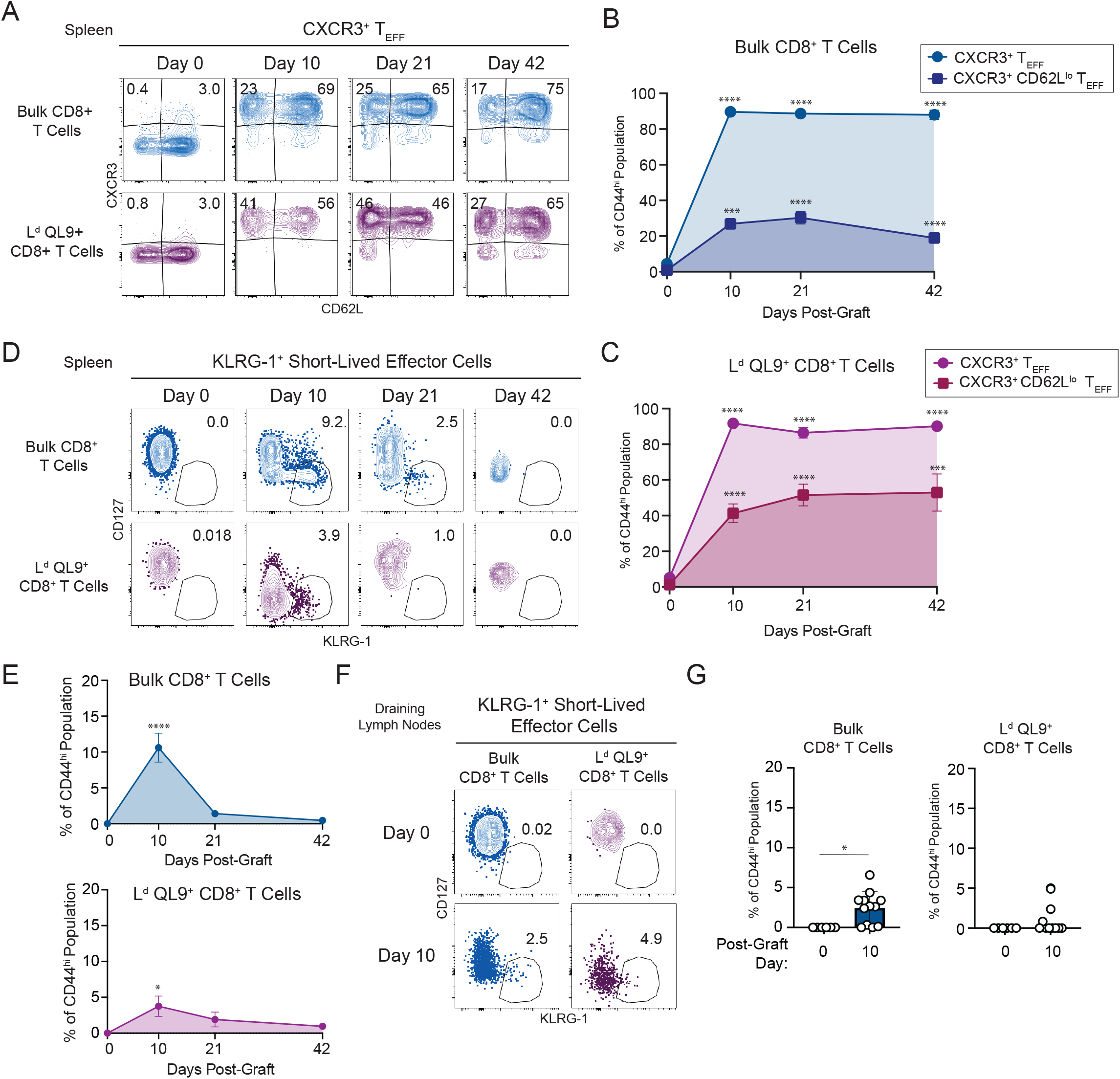
Post-graft effector CD8^+^ T cells express activation markers but poorly express KLRG-1. C57Bl/6 mice were grafted with H-2^d^ Balb/c skin. (A) Representative flow cytometry plots and summary data depicting the frequency of CXCR3^+^ and CD62L^lo^CXCR3^+^ T_EFF_ among (B) bulk CD8^+^ T cells and (C) L^d^ QL9^+^ CD8^+^ T cells at the indicated timepoints post-graft. (D) Representative flow cytometry plots and (E) summary data depicting the frequency of CD127^lo^KLRG1^+^ cells among bulk CD8^+^ T cells and L^d^ QL9^+^ T cells at the indicated timepoints in the spleen. (F) Representative flow cytometry plots and (G) summary data depicting the frequency of CD127^lo^KLRG-1^+^ cells among bulk CD8^+^ T cells and L^d^ QL9^+^ CD8^+^ T cells at day 0 or day 10 in the graft draining lymph nodes. Each data point depicts an individual mouse. Summary data depicts pooled results from 2-3 independent experiments (n=6-12 mice/group). Error bars depict SEM.

The early CD8^+^ T cell response is often defined by the reciprocal expression of the CD127 (IL-7R α) and KLRG-1 receptors. In the first 1-2 weeks after infection, a significant portion (50-70%) of T_EFF_ transiently downregulate CD127 and become KLRG-1^+^, and these cells are short-lived and display potent effector functions(Herndler-Brandstetter et al., 2018b; Joshi et al., 2007; Milner et al., 2020). We assessed the expression of CD127 and KLRG-1 among L^d^ QL9^+^ T_EFF_ after grafting in both the spleen and draining lymph nodes. In response to a Balb/c skin graft, a low frequency of CD127^lo^KLRG1^+^ T_EFF_ cells were induced among L^d^ QL9^+^ and bulk CD8^+^ T cells in the spleen, with a peak frequency at day 10 that declined precipitously at day 21 and 42 **(Figure 4D-E)**. In the draining lymph node, a similarly low frequency of KLRG-1^+^ T_EFF_ were found **(Figure 4F-G)**. Thus, these data show that among graft-specific L^d^ QL9^+^ cells after transplantation, KLRG-1 expression does not define an abundant population of acutely responding T_EFF_.

### Expression of the activated CD43 glycoform is increased on a population of CD8^+^ T_EFF_ after grafting

To better characterize the phenotypes of the CD8^+^ T_EFF_ cells after grafting, we analyzed a high-parameter flow cytometry panel with canonical costimulatory and activation using dimensionality reduction analysis. We evaluated L^d^ QL9^+^ and bulk CD8^+^ T cells on day 0, 10, 21, and 42 after Balb/c skin grafting by clustering these populations with FlowSOM and ConsensusClusterPlus and visualized with uniform manifold approximation and projection (UMAP) dimensionality reduction **(Figure 5A-C)**. We evaluated the frequency of clusters and found four clusters that were enriched on day 10 relative to the other timepoints, and three clusters that were enriched on both day 21 and 42 **(Figure 5B-C)**. We further evaluated day 10 enriched clusters, as these fit the profile of a short-lived effector population. All four Day 10 Clusters were CD44^hi^, Ki67^+^, CXCR3^hi^ and KLRG-1^-^ **(Figure 5D)**. Interestingly, these clusters varied most in expression of CD62L and the activated glycoform of CD43 (identified by the antibody clone 1B11). While Day 10 Clusters 1 and 2 were CD62L^hi^CD43^-^, Cluster 3 was CD62L^lo^CD43^-^, and Cluster 4 was CD62L^lo^CD43^+^ (**Figure 5D)**. Thus, these two surface receptors appear to represent dynamic populations within the T_EFF_ pool after grafting. In order to evaluate the expression of CD62L and CD43 after grafting, we performed manual gating on L^d^ QL9^+^ and bulk CD8^+^ T_EFF_. We found that the frequency of CD62L^lo^CD43^+^ T_EFF_ increased at day 10 and 21 and were decreased at a memory day 42 timepoint **(Figure 5E)**. Thus, activated CD43 expression defines a population of CD62L^lo^ T_EFF_ that appears acutely after grafting and declines over time, fitting the kinetic profile of an acutely responding CD8^+^ T_EFF_ population.

**Figure 5.**
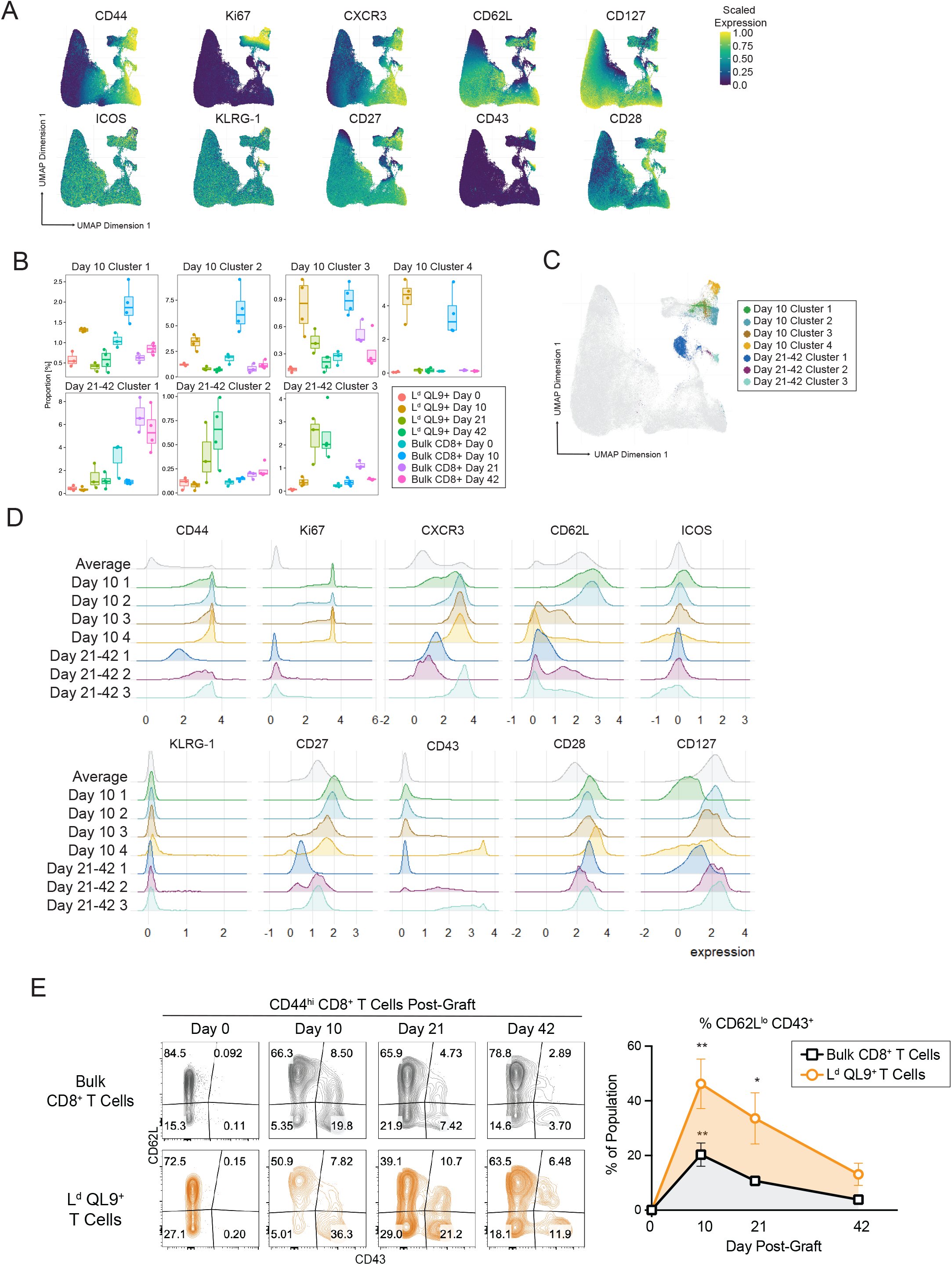
CD8^+^ T cells differentiate into CD43^+^ T effector populations that peak early after grafting. C57Bl/6 mice were grafted with H-2^d^ Balb/c skin and the frequency of L^d^ QL9^+^ and bulk CD8^+^ T cells were assessed in the spleen at the indicated timepoints. (A) UMAP of costimulation and activation markers from day 0, 10, 21, and 42 post-graft CD8^+^ T cell populations. (B) Box plots of the frequency of clusters elevated at day 10 or day 21/42. (C) UMAP depicting day 10 and day 21/42 enriched clusters. (D) Histograms depicting expression of individual markers in day 10 and day 21/42 enriched clusters. (E) Expression of CD43 and CD62L on CD8^+^ T cell populations at day 0, 10, 21, and 42 post-graft. Each data point depicts an individual mouse. Summary data depicts pooled results from (A-D) one (n=4 mice/group), (E) 2-3 independent experiments (n=5-10 mice/group). Statistics performed by (B-C, E) 1-way ANOVA with Dunnett’s multiple comparison’s test or (G) unpaired Student’s t-test. Error bars depict SEM. *p<0.05, **p<0.01, ***p<0.001, ****p<0.0001.

### CD43^+^ CD8^+^ T_EFF_ display potent effector functions relative to CD43^-^ T_EFF_ populations

CD43 is a transmembrane receptor that has been shown to impact intracellular signaling cascades, apoptosis, and T cell trafficking. However, the role of activated CD43 expression on CD8^+^ T cells has not been evaluated. Thus, we next sought to assess whether CD43^+^ T_EFF_ displayed competent effector functions relative to CD43^-^ T_EFF_ populations. We grafted C57Bl/6 mice with Balb/c skin and assessed CD8^+^ T cells during the peak effector response (day 10-14; **Figure 6A**). We found that among the L^d^ QL9^+^ cells, CD43^+^ T_EFF_ expressed significantly higher levels of both Granzyme B and T-Bet as compared with CD62L^hi^ or CD62L^lo^ CD43^-^ T_EFF_ populations **(Figure 6B-C)**. We assessed the capacity of these populations for cytokine production by stimulating CD45.1^+^ T_EFF_ *ex vivo* with Balb/c splenocytes, and found that nearly all of the IFN-γ and TNF-α was produced by CD43^+^ T_EFF_, and that the majority of these cells were double producers **(Figure 6D)**.

**Figure 6.**
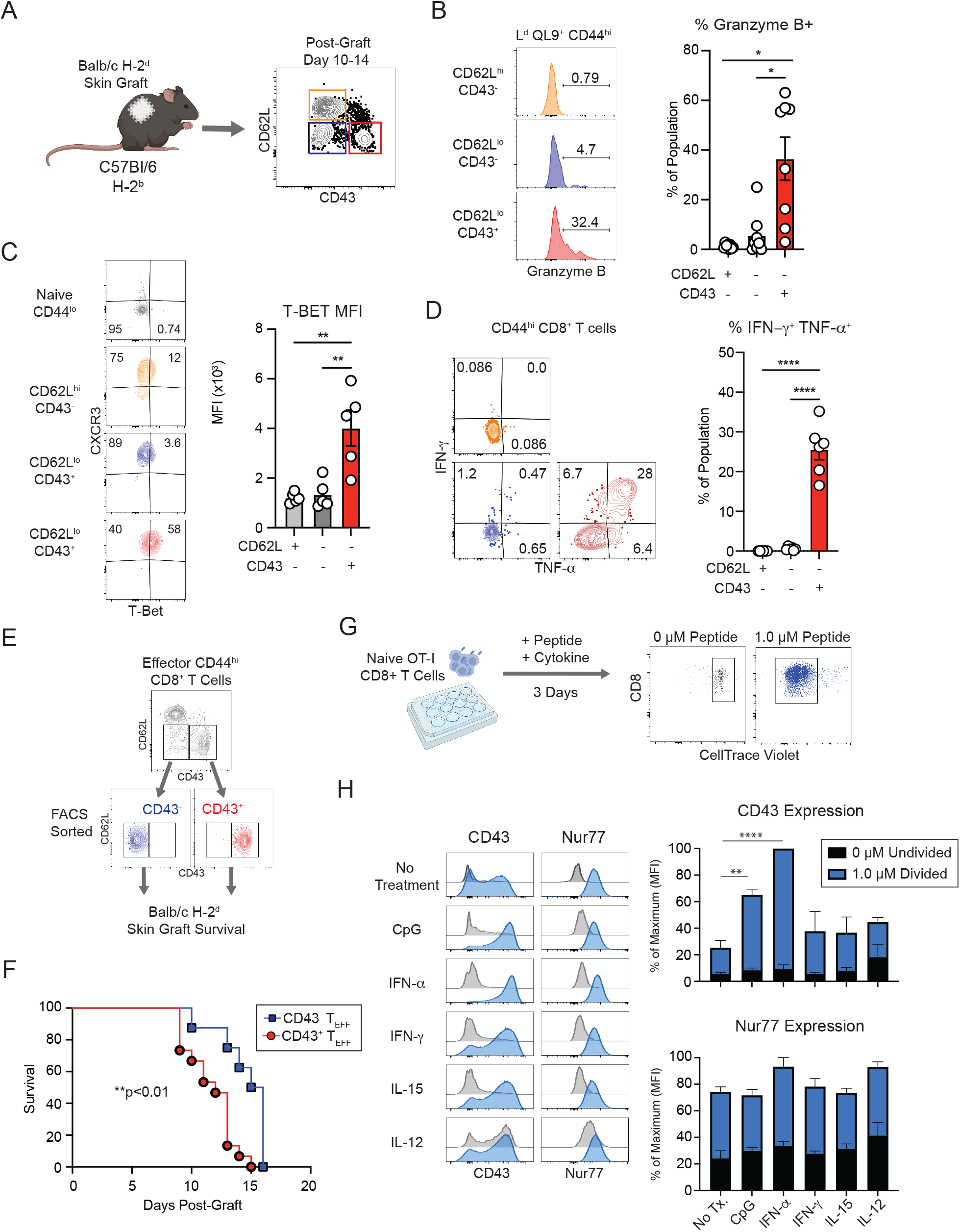
CD43^+^ T_EFF_ display potent effector function and mediate graft rejection. **(A)** C57Bl/6 mice were grafted with H-2^d^ Balb/c skin and L^d^ QL9^+^ and bulk CD8^+^ T cells were assessed in the spleen at day 10-14 post-graft. Representative flow cytometry plots and summary data of (B) Granzyme B^+^ cells and (C) T-BET^+^ cells in the indicated CD8^+^ T_EFF_ populations. (D) Post-graft effectors were stimulated *in vitro* for 5 hours with Balb/c splenocytes and the frequency of IFN-ψ and TNF-α producing cells was assessed. (E) CD45.1^+^ CD8^+^ T cells were sorted into CD62L^lo^ CD43^+^ and CD43^-^ populations, and 9×10^4^ cells/mouse were transferred into congenic hosts that were grafted with Balb/c skin grafts and assessed for survival. (F) Graft survival analysis of C57Bl/6 hosts transferred with CD45.1^+^ CD8^+^ T cell populations described in (E). (G) OT-I CD8^+^ T cells were labeled with CellTrace Violet and stimulated *in vitro* for 3 days in the presence of 0 µM (unstimulated) or 1 µM OVA peptide and the indicated cytokine. (H) Histograms depicting the expression of CD43 and Nur77 on undivided unstimulated or 1 µM peptide divided populations. For each individual experiment, the MFI was normalized to the maximum value of CD43 or Nur77. Each data point depicts an individual mouse. Summary data depicts pooled results from (B-D) 2 independent experiments (n=5-8 mice/group), (F) 2 independent experiments (n=7-15/group), (H) 3 independent experiments with an individual mouse. Statistical analyses performed by (B-D) 1-way ANOVA with Tukey’s multiple comparisons test, (F) Log-rank (Mantel-Cox) test, (H) 2-way ANOVA with Dunnett’s multiple comparisons test. Error bars depict SEM. *p<0.05, **p<0.01, ****p<0.0001.

We wanted to assess whether the effector phenotype corresponded with an enhanced potency to mediate graft rejection *in vivo*. To isolate the impact of CD43 on T_EFF_, we FACS-sorted CD44^hi^ CD8^+^ T cells into CD43^+^ and CD43^-^ T_EFF_ (CD62L^lo^) on day 10-14 and adoptively transferred 9×10^4^ cells into C57Bl/6 mice who were subsequently grafted with Balb/c skin. We found that the CD43^+^ T_EFF_ group rejected skin grafts with significantly faster kinetics than CD43^-^ T_EFF_ group **(Figure 6E-F)**, demonstrating the impact of CD43^+^ T_EFF_ to mediate potent graft rejection. Overall, these data demonstrate that CD43^+^ T_EFF_ are a potent population of effector CD8^+^ T cells *in vivo*.

### Activated CD43 expression is induced by antigen and Type I interferon

Our data show that activated CD43 is induced after grafting *in vivo*, which entails exposure to both allogeneic antigen and inflammatory cytokines. We questioned whether inflammatory cytokines alone drive the expression of activated CD43 during T cell priming. Multiple studies have demonstrated that pathogenic inflammation on the development and function of CD8^+^ T cell populations(Joshi et al., 2007; Richer et al., 2015). We activated OT-I T cells *in vitro* in the presence of individual inflammatory cytokines **(Figure 6G)**. We used CellTrace Violet to assess the proliferation status of cells in culture. We included expression of Nur77, an orphan nuclear receptor whose expression level reflects the cumulative strength of antigen stimulation(Krummey et al., 2016; Moran et al., 2011), as a control for the overall level of activation of the cells in the presence of cytokines. We found that antigen stimulation alone induced CD43 and Nur77 expression **(Figure 6H)**. The TLR9 agonist CpG, which has been shown to induce CD43 (1B11) expression(Nolz and Harty, 2014), increased CD43 expression further than antigen stimulation alone. Among inflammatory cytokines, we found that only IFN-α induced CD43 expression, while IFN-γ, IL-15, and IL-12 did not have an effect versus antigen stimulation alone **(Figure 6H)**. The expression of Nur77 was not significantly impacted by the provision of any inflammatory cytokine. These results demonstrate that CD43 (1B11) expression is enhanced by antigen and can be augmented by Type I IFN mediated signaling.

### Costimulation blockade with CTLA-4 Ig does not inhibit CD8^+^ T cells

Costimulation blockade with CTLA-4 Ig or a biochemical derivative is used to treat autoimmune disease and to prevent transplant rejection in kidney transplant patients. In transplant patients, CTLA-4 Ig is associated with acute T cell mediated rejection, and pre-clinical studies have shown that CD8^+^ T cells are relatively resistant to CTLA-4 Ig (Crepeau and Ford, 2017; Ford et al., 2007; Liu et al., 2014). We next evaluated the impact of CTLA-4 Ig on CD43^+^ T_EFF_ relative to other T_EFF_. We treated mice with CTLA-4 Ig in comparison to a control untreated group and analyzed CD8^+^ T cells on day 10 after grafting with Balb/c skin grafts **(Figure 7A)**. Consistent with prior studies, we found that CTLA-4 Ig reduced the proliferation of effector CD4^+^ T cells but did not restrain CD8^+^ effector T cell populations **(Figure S2)**.

**Figure 7.**
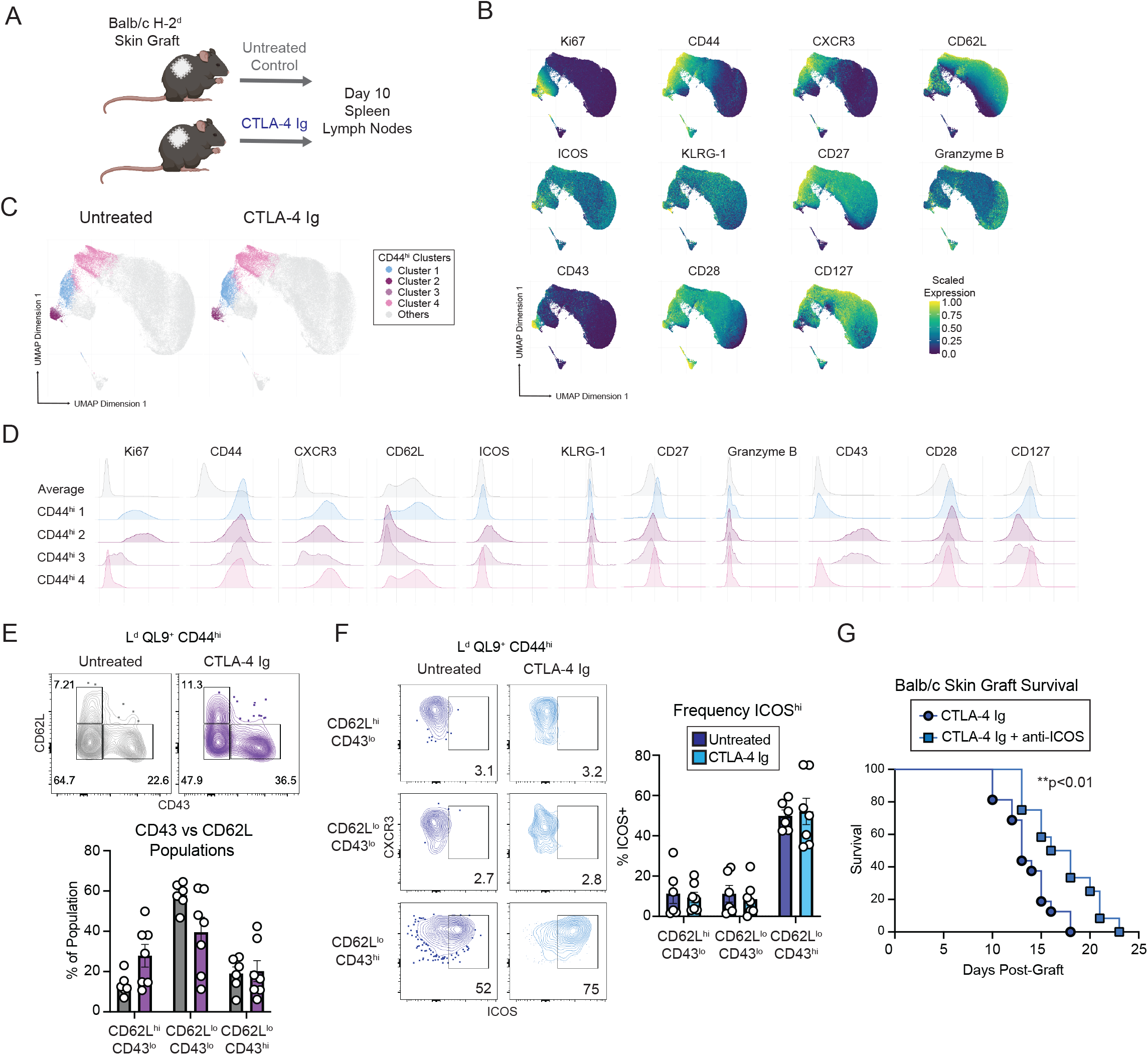
ICOS expression is induced on CD43+ T_EFF_ and ICOS blockade prolongs graft survival. (A) Experimental schematic depicting the treatment of C57Bl/6 mice grafted with H-2^d^ Balb/c skin and treated with CTLA-4 Ig. (B) UMAP expression analysis of day 10 post-graft L^d^ QL9^+^ and bulk CD8^+^ T cells in the spleen in mice untreated or treated with CTLA-4 Ig. (C) UMAP depicting four CD44^hi^ clusters. (D) Histograms of proteins used in UMAP analysis for four CD44^hi^ clusters and the average of all samples. (E) Representative flow cytometry plots and summary data depicting the frequency of CD62L and CD43-expressing effector populations in the spleen on day 10 post-graft in mice. (F) Expression of ICOS on CD8^+^ T_EFF_ populations in CTLA-4 Ig and control mice. (G) Graft survival of C57Bl/6 mice treated with CTLA-4 Ig or CTLA-4 Ig with anti-ICOS and grafted with a Balb/c skin graft. Each data point depicts an individual mouse. Summary data depicts pooled results from (E-F) 2 independent experiments (n=6-7 mice/group), (G) 3 independent experiments (n=12-16/group). Statistical analyses performed by (E-F) 2-way ANOVA with Tukey’s multiple comparisons test, (G) Log-rank (Mantel-Cox) test. Error bars depict SEM. **p<0.01

In order to evaluate the phenotype of CD8^+^ T cells in the presence of CTLA-4 Ig, we performed UMAP and clustering analysis of L^d^ QL9^+^ and bulk CD8^+^ T cells in treated and control mice **(Figure 7B)**. We found four clusters that were CD44^hi^ T_EFF_ and evaluated the expression of costimulation receptors, activation markers, and functional markers **(Figure 7B-D)**. CD44^hi^ Clusters 2 and 3 had high CD43 (1B11) expression, and were also CD62L^lo^, CXCR3^hi^, and Ki67^+^. Interestingly, both had high expression of the costimulation receptor ICOS **(Figure 7D)**. Using manual gating, we evaluated whether the frequency of CD62L and CD43 T_EFF_ was impacted by CTLA-4 Ig treatment, and found that all three populations were maintained at similar frequencies at day 10 between groups **(Figure 7E)**. We next evaluated ICOS expression on CD43 vs CD62L T_EFF_, and found that ICOS expression was almost exclusively found on CD62L^lo^CD43^+^ T_EFF_ **(Figure 7F)**. In order to assess whether ICOS costimulation provisioned functional signals in the context of graft rejection, we assessed the impact of ICOS blockade in combination with CTLA-4 Ig treatment. We found that anti-ICOS prolonged graft survival versus CTLA-4 Ig alone **(Figure 7G)**. Thus, ICOS represents a functional costimulation target that can be used to inhibit CTLA-4 Ig resistant CD8^+^ T cells.

## DISCUSSION

CD8^+^ T cells are thought to respond to allogeneic antigen primarily through the direct antigen presentation pathway, in which donor MHC complexed with ubiquitously expressed self peptides bind to host CD8^+^ T cells (Harper et al., 2015; Siu et al., 2018). Burrack et al recently used MHC Class II tetramers to show that CD4^+^ T cells can respond to directly presented allogeneic antigen in a mouse model of skin grafting (Burrack et al., 2018), but allogeneic CD8^+^ T cells have not been studied using a similar approaches (Young et al., 2017). In this study, we build upon several elegant studies by Eisen and colleagues to study the CD8^+^ T cell response directed against the H-2L^d^ complexed with the self-peptide QL9. While this early work defined the biochemistry of the 2C TCR binding with L^d^ QL9 *in vitro* (Chen et al., 2003; Sykulev et al., 1994a, 1994b, 1996; Udaka et al., 1992, 1996), the *in vivo* immune CD8+ T cell response directed at L^d^ QL9 has not been evaluated.

Thus, we sought to use MHC Class I tetramer enrichment approach to study L^d^ QL9-specific CD8^+^ T cells. We found that at the peak of the immune response, the majority of L^d^ QL9^+^ T cells become CD44^hi^ and nearly all CD44^hi^ L^d^ QL9^+^ T cells are Ki67^+^, providing strong evidence that the tetramer binding T cells recognize endogenously presented L^d^ QL9 on graft tissue. While the QL9 peptide is a single amino acid extension of the parent 2C peptide, prior studies investigating the peptides recognized by the 2C TCR speculated that the QL9 was not detected on donor antigen-presenting cells due to technical limitations of peptide sequencing in acidic conditions (Chen et al., 2003; Sykulev et al., 1994).

We found that the precursor frequency for L^d^ QL9^+^ T cells is elevated compared with K^b^ OVA and D^b^ LCMV gp33, as well as the published frequencies of other pathogen-specific antigens (Haluszczak et al., 2009; Jenkins and Moon, 2012; Kotturi et al., 2008; Obar et al., 2008). Prior studies estimating the precursor frequency of alloreactive T cells have relied on a wide variety of techniques, including *in vitro* stimulation for proliferation or cytokine production, and have relied on memory-containing T cell populations and/or expanded T cell lines (Amir et al., 2010; Felix et al., 2007; Heeger et al., 1999; Macedo et al., 2009; Suchin et al., 2001). Thus, our findings using an MHC tetramer to directly visualize epitope-specific CD8^+^ T cells significantly strengthens the knowledge about the precursor frequency of alloimmunity gained from these prior studies.

Our results provide a model in which alloreactive CD8^+^ T cells exist at a high precursor frequency but undergo relatively weak clonal expansion. Consistent with this, *TCRB* sequencing of the L^d^ QL9-specific CD8+ T cell response reveals a highly diverse response at the clonal level. While we identified biochemically similar group of convergent meta-clonotypes that expanded after grafting, the anti-graft CD8^+^ T cell response remained diverse between individual animals. Interestingly, while we observed that the meta-clonotypes were present in graft tissue, the graft-infiltrating response was not dominated by the convergent clones at day 10 post-graft. Future studies are needed to gain a better understanding of the clonal properties and kinetics of the CD8^+^ T cell infiltration into graft.

In contrast to pathogen-specific CD8^+^ T cell responses at acute timepoints, we found that very few effector CD8^+^ T cells expressed KLRG-1 after grafting. This highlights that the allogeneic priming environment has phenotypic consequences in shaping CD8^+^ T cell responses, and led us to carefully evaluate the post-graft CD8^+^ T cell programming. Using a high-parameter flow cytometry panel to evaluate activation, costimulation, and proliferation markers, we found that a subset of L^d^ QL9^+^ T_EFF_ express the activated glycoform of CD43 (1B11 clone). The frequency CD43^+^ T_EFF_ increased acutely after transplant and these cells were potently cytotoxic relative to CD43^-^ T_EFF_ populations.

CD43 is a transmembrane receptor with recognized intracellular signaling and adhesion functions (Mody et al., 2007; Onami et al., 2002; Sperling et al., 1995). CD43 is uniformly high on T cells, but a larger activated glycoform, defined by the 1B11 epitope, is selectively expressed in certain contexts (Clark and Baum, 2012). Studies evaluating the role of CD43 in disease models have found varying and sometimes conflicting consequences. CD43 deficient CD8^+^ T cells have been found to undergo a decreased rate of apoptosis during the contraction phase of the response to a virus (Onami et al., 2002), but including increased levels of apoptosis in a model of sepsis (Fay et al., 2018). Multiple studies have shown that CD43 is critical for trafficking into peripheral tissues (Cannon et al., 2011; Ford and Evavold, 2006; Ford et al., 2003; Mody et al., 2007; Onami et al., 2002). Of note, the majority of studies have evaluated the function of CD43 sufficient versus strains, less is known about the functional impact of CD43 1B11 glycoform expression on T cells. Future studies will be needed to evaluate the importance of CD43 1B11 expression on the function of T_EFF_ in the context of transplant, including the ability to traffic within secondary lymphoid tissue and into allografts.

We evaluated antigen-specific T_EFF_ in the presence of costimulation blockade with CTLA-4 Ig, and found that the costimulatory receptor ICOS is selectively expressed on CD43^+^ T_EFF_. Blockade of ICOS prolonged graft survival, indicating that ICOS plays a functional role in the presence of CTLA-4 Ig treatment. ICOS has been implicated as an important costimulatory receptor in multiple T cell subsets, and was recently shown to be important for the formation of resident memory CD8^+^ T cells (Peng et al., 2022). ICOS is attractive as a therapeutic target because it is selectively and transiently upregulated on CD43^+^ T_EFF_, and thus represents a potentially selective way to inhibit allogeneic CD8^+^ T cells during rejection. Currently, ICOS blocking agents are in pre-clinical development for multiple diseases (Adom et al., 2020; Solinas et al., 2020).

Overall, by using a novel MHC tetramer-based approach, this study provides new insight into the characteristics of the alloimmune CD8^+^ T cell response, including the identification of a potent T_EFF_ phenotype. Future studies are needed to investigate the translational potential of these findings to inhibit graft-specific CD8^+^ T cells in transplant patients.

## ACKNOWLEDGEMENTS

We thank members of the Ford Lab and the Immunology Division of the Department of Pathology at Johns Hopkins for helpful discussions. We appreciate Hao Zhang and the Bloomberg School of Public Health Flow Cytometry and Cell Sorting Core for excellent cell sorting, and the NIH Tetramer Core Facility for tetramer reagents and technical advice. This work was supported by K99/R00 NIAID AI46271 (S.M.K) and the Transplant Immunology Research Network Fellowship Grant (S.M.K.).

## AUTHOR CONTRIBUTIONS

Conceptualization, S.M.K., H.B.L., G.A.C., S.J.; methodology, S.M.K., G.A.C., H.B.L., S.J.; formal analysis, S.M.K., G.A.C., S.A.J., H.B.L.; investigation, S.M.K., G.A.C., M.A.K., S.J., F.I.I., K.P.T., L.A.O.; writing, S.M.K., G.A.C., S.J.; visualization, S.M.K., G.A.C.; supervision, S.M.K., H.B.L.

## METHODS

### Mice

C57BL/6J and C57BL/6J Ly5.2-Cr (CD45.1) strains were obtained from The Jackson Laboratory (Bar Harbor, Maine) at 6-8 weeks of age. Balb/cJ, C57Bl/6-Tg(TcraTcrb)1100Mjb/J (OT-I), and RAG1^tm1Mom^ (RAG1 knockout) were obtained from The Jackson Laboratory and bred at Emory University or Johns Hopkins University. All transplant experiments were conducted in age-matched hosts randomly assigned to experimental groups. This study was conducted in accordance with the recommendations in the Guide for the Care and Use of Laboratory Animals. The protocol was approved by the Animal Care and Use Committee at Emory University and Johns Hopkins University. Animals were housed in specific pathogen-free animal facilities at Emory University and Johns Hopkins University.

### Skin Grafting

Full-thickness tail and ear skins were transplanted onto the dorsal thorax of recipient mice and secured with adhesive bandages as previously described (Trambley et al., 1999). In some experiments, mice were treated with 250 ug/dose of CTLA-4 Ig (BioXCell) and anti-ICOS (BioXCell) via i.p. injection on day 0, 2, 4, and 6.

### MHC Tetramers and CD8^+^ T cell Enrichment

MHC tetramers with human β2-microglobulin specific for QL9 (H-2L^d^ QLSPFPFDL), LCMV gp33 (H-2D^b^ KAVYFATC), and OVA (H-2K^b^ SIINFEKL) were obtained from the NIH Tetramer Core facility. Tetramer enrichment was performed based on published protocols (Moon et al., 2009). Briefly, single cell suspensions of splenocytes or lymphocytes were incubated with MHC tetramer (PE or APC), followed by incubation with anti-PE or anti-APC paramagnetic microbeads (Miltenyi), and enrichment over an LS column (Miltenyi). The column flow through was collected as the unbound fraction containing bulk CD8^+^ T cells and the column bound fraction enriched for tetramer binding cells were stained for surface and intracellular antigens and analyzed by flow cytometry. Absolute cell counts were performed using CountBrite beads (BD Biosciences) or Cytek Aurora.

### Antibodies and Flow Cytometry

Flow cytometry and magnetic enrichment were performed in 1X PBS (without Ca^2+^/Mg^2+^, pH 7.2), 0.25% BSA, 2 mM EDTA, and 0.09% azide. Single cell suspensions were incubated in 96-well U-bottom plates with the following antibodies for 30-60 min at room temperature: CD4, CD8, CD11c, F4/80, CD19, CD27, CD28, CD127, KLRG-1, CD43, CXCR3, CD62L, ICOS, CD44, CD62L. Intracellular staining was performed using the Transcription Factor Staining Kit (eBiosciences) and the following antigens: Ki67, Nur77, Granzyme B, T-Bet, IFN-γ, TNF-α. Dead cells were excluded with Live/Dead Aqua (Invitrogen), Live/Dead Near IR (Biolegend), or propidium iodide (Invitrogen) according to manufacturer’s instructions.

### *Ex vivo* Allogeneic Stimulation for Cytokine Production

Splenocytes and draining lymph node cells from CD45.1^+^ mice grafted with Balb/c skin grafts 14 days prior were processed to single cell suspension and pooled. 1.5×10^6^ CD45.1 cells were incubated with 2×10^6^ Balb/c splenocytes in the presence of 1 mg/mL GolgiStop (Invitrogen) for 4-5 h at 37 C. Cells were collected and gated on CD45.1^+^ CD8^+^ T cells for intracellular cytokine analysis.

### Flow cytometric cell sorting and adoptive transfer

Splenocytes and draining lymph nodes cells from CD45.1^+^ mice grafted with Balb/c skin grafts 10-14 days prior were stained to identify effector CD62L^lo^CD43^+^ and CD62L^lo^CD43^-^ CD8^+^ T cells. Cells were sorted on a MoFlo XDP. 9×10^4^ purified cells were transferred via tail vein into C57BL/6J mice. Mice were grafted the following day with Balb/c skin and monitored for graft survival.

### *In vitro* OT-I Stimulation

Naïve OT-I splenocytes were stained with CellTrace Violet (Invitrogen) according to the manufacturer’s instructions and plated in 24 well tissue culture plates at 3×10^6^/well in in the presence of OT-I peptide (GenScript) and the following cytokines: CpG (3.125 µg/mL, Invivogen), IFN-α (Biolegend, 1.1 µg/mL), IL-12 (Biolegend, 10 ng/mL), IL-15 (Biolegend, 10 ng/mL), IFN-γ (Biolegend, 100 ng/mL). Cells were incubated in Complete R10 media comprised of RPMI 1640 with L-glutamine (Corning), 10% FBS, 100 mM HEPES, 500 uM ϕ-mercaptoethanol, 100 U/mL penicillin/stremptomycin at 37 C (5% CO_2_) for 72 h prior to analysis by flow cytometry.

### *TCRB* Sequencing (FR3AK-Seq) and Analysis

CD8^+^ T cell populations from splenocytes and draining lymph nodes were flow cytometrically sorted from naïve or post-Balb/c skin graft mice as described above. RNA was isolated using the RNEasy Plus Mini kit (Qiagen) according to manufacturer instructions. cDNA was generated using the Superscript IV First-Strand Synthesis System kit per manufacturer protocol instructions. A normalized input of 200ng of RNA was used for each sample, or if the concentration was too low to reach 200ng, the maximum amount of RNA that could be input into the reaction was used. RNA corresponding to 1000 cells of the EL4 cell-line was used as a spike-in in all samples. cDNA samples were cleaned-up and concentrated using the Zymo Research ZR-96 DNA Clean and Concentrator kit (cat# D4023). 20 cycles of PCR1 were performed using the KAPA2G Fast Multiplex Mix and primers designed to encompass all mouse TCRBV alleles. 20 cycles of PCR2 was performed for sample barcoding. Sequencing was performed on an Illumina NextSeq. CDR3s were identified and quantified using MiXCR software (v 3.0.13), and the EL4 spike-in was used to normalize MiXCR clone counts.

Assembled clonotypes were imported and analyzed in R (4.1.1) via the immunarch (0.6.7) package. Clonotypes were categorized for expansion based on the relative frequency it represented in that sample using raw clone counts as follows: >1% hyper-expanded, 1%-0.1% large, 0.1%-0.01% medium, <0.01% small. Graphics were generated with ggplot2 (3.3.5). For any analysis, that required a defined v or j gene and there was an ambiguous result, the top hit was taken. Metaclonotypes were generated in python (3.8.12) using tcrdist3 (0.2.2). Data were imported using pandas (1.3.4). The enriched clonotypes consisted of sorted spleen cells, that were tetramer bound, and CD44^hi^. The background set was generated from naïve mouse blood samples (n=14). A random sample of 280,000 of these clonotypes were generated with tcrsampler (0.1.9), and a v-j matched background of 280,000 clonotypes was generated with olga (1.2.4).

### Flow Cytometry Analysis

In all analysis, CD8^+^ T cells were identified as singlet events, Live/Dead stain^-^, dump gate^-^ (CD11c, F4/80, NK1.1, CD19), CD4^-^, CD8^+^ events. Manual gating was performed in Flowjo 10. Tetramer and bulk CD8^+^ T_EFF_ are defined as CD44^hi^CD8^+^ T cells. For dimensionality reduction analysis, manually gated unbound CD8^+^ T cells and bound fraction L^d^ QL9^+^ CD8^+^ T cells events were imported into R (4.1.1) through CytoML (2.40), flowWorkspace (4.4.0), and flowCore (2.4.0). Markers were transformed by hyperbolic arcsine with a cofactor of either 1000 or 6000 depending on the marker. The data were further analyzed using CATALYST (1.16.2) with FlowSOM (2.0.0) clustering and ConsensusClusterPlus (1.56.0) meta clustering. UMAP dimensionality reductions were generated with scatter (1.20.1). Visualizations were generated in ggplot2 (3.3.5) and ComplexHeatmap (2.8.0).

### Statistical Analysis

All data points represent individual mice, and where individual data points are not depicted the value of n is provided in the corresponding figure legend. For analysis of absolute numbers and expression levels, paired or unpaired Student’s t-tests (two-tailed) were performed between two groups; one-way or two-way ANOVA with multiple comparison tests were used to compare multiple groups. Log-rank (Mantel-cox) test was used to evaluate graft survival between groups. In vitro CD43 and Nur77 expression data was normalized to the maximum MFI obtained for each marker in a given experiment. Error bars represent standard error measurements (SEM). Statistics were performed using GraphPad Prism 9. Significance was determined as *p<0.05, **p<0.01, ***p<0.001, ****p<0.0001.

**Table S1.**
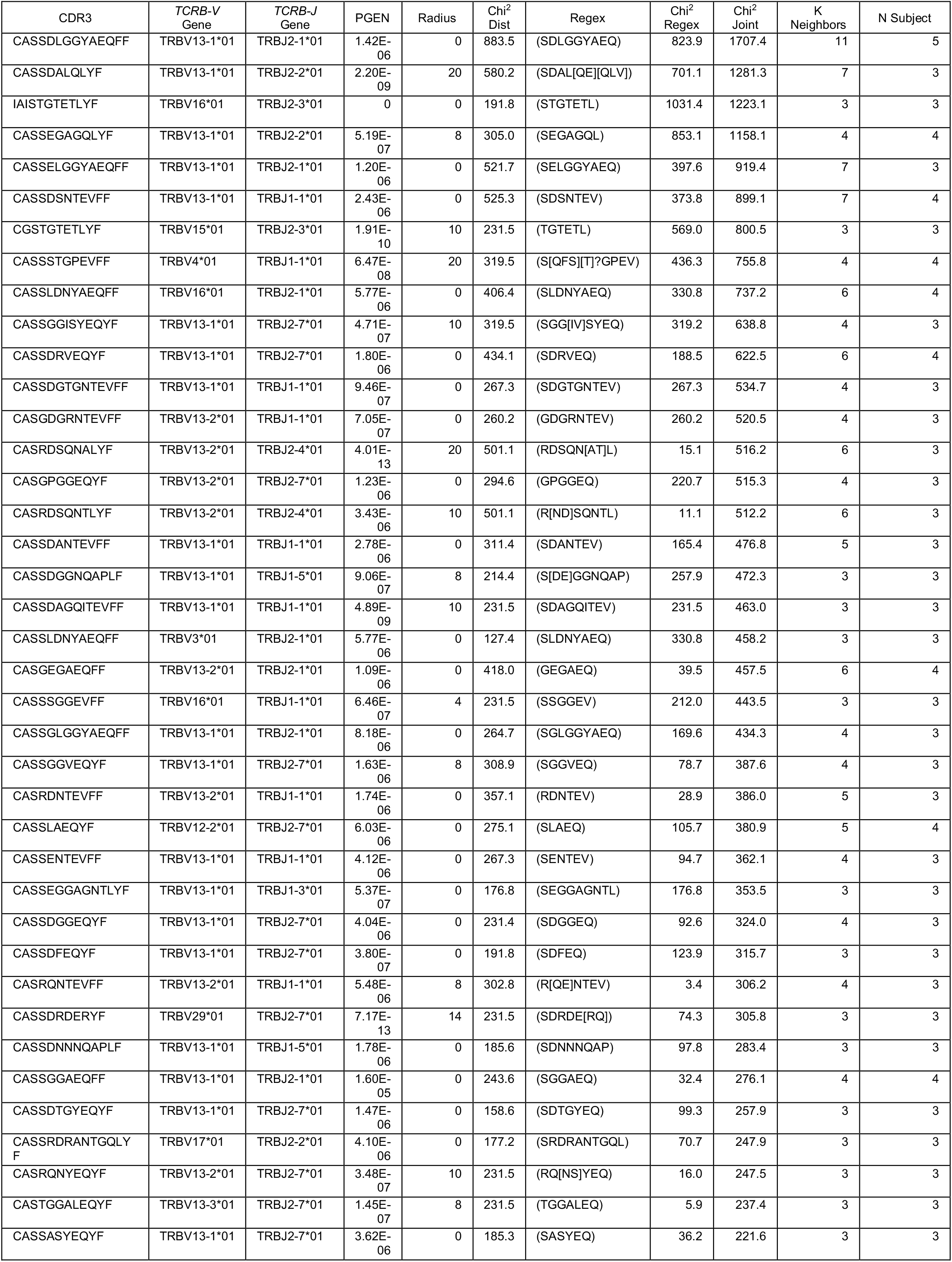
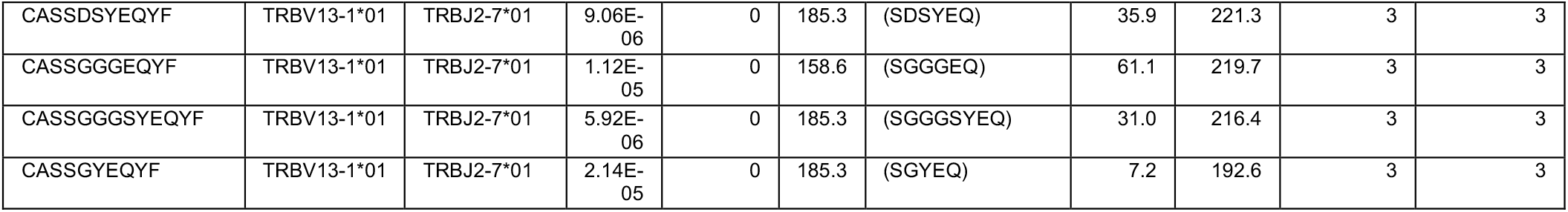
Meta-clonotypes identified in naïve and effector L^d^ QL9^+^ effector CD8^+^ T cells. *TCRB* CDR3 sequences from sorted L^d^ QL9+ and bulk T cell populations were analyzed using the tcrDist package. Forty-three meta-clonotypes were identified that met criteria of being enriched over background populations and present in more than two samples.

**Figure S1.**
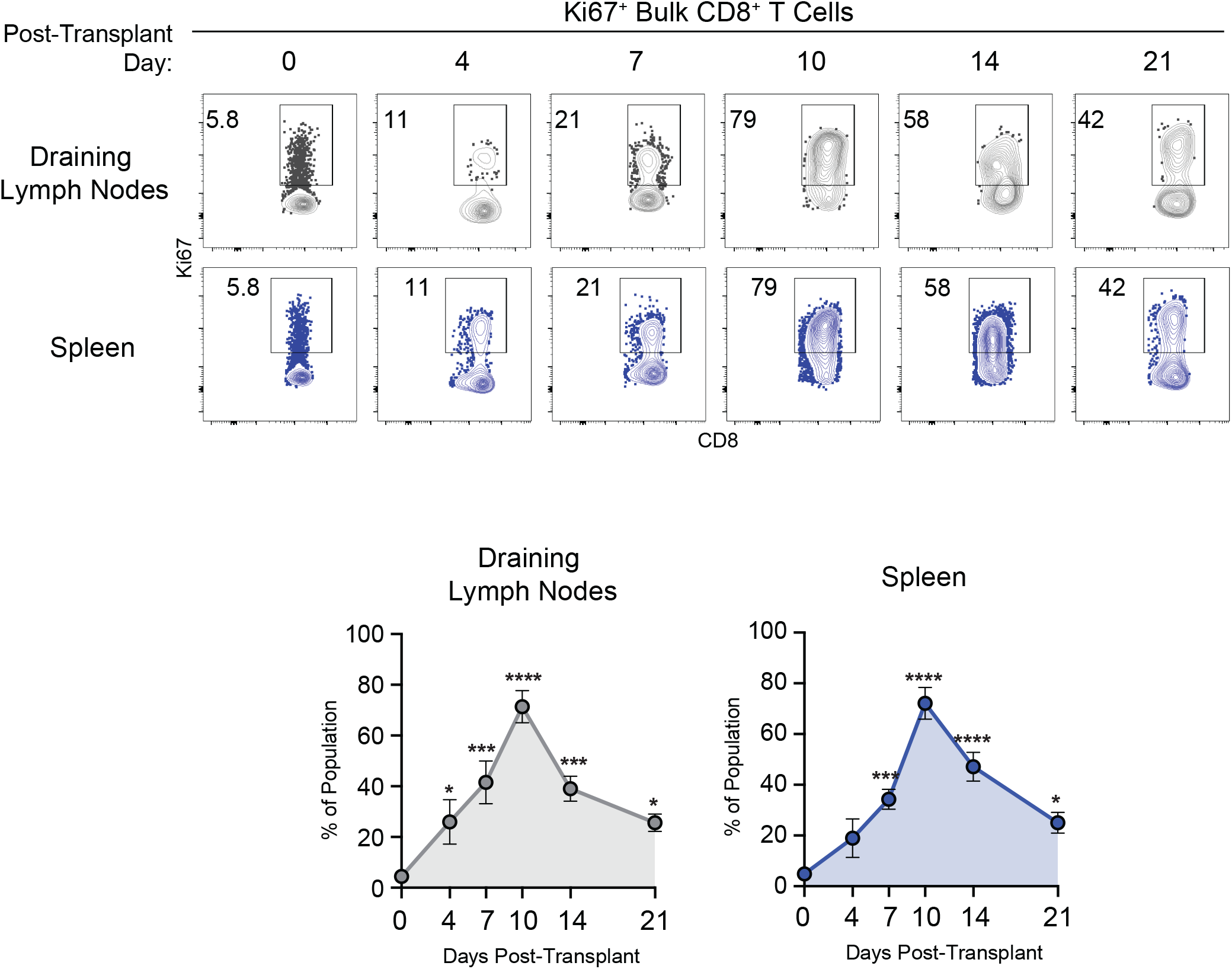
Bulk CD8^+^ T cells proliferate in the spleen and draining lymph nodes after grafting. Related to Figure 2. C57Bl/6 mice were grafted with H-2^d^ Balb/c skin. Representative flow cytometry plots and summary data of Ki67 staining among CD44^hi^ CD8^+^ T cell populations in the draining lymph nodes and spleen. Each data point depicts an individual mouse. Summary data depicts pooled results from 2-3 independent experiments (6-9/group). Statistical analysis performed using one-way ANOVA with Dunnett’s multiple comparisons test. Error bars depict SEM. *p<0.05, **p<0.01, ***p<0.001, ****p<0.0001.

**Figure S2.**
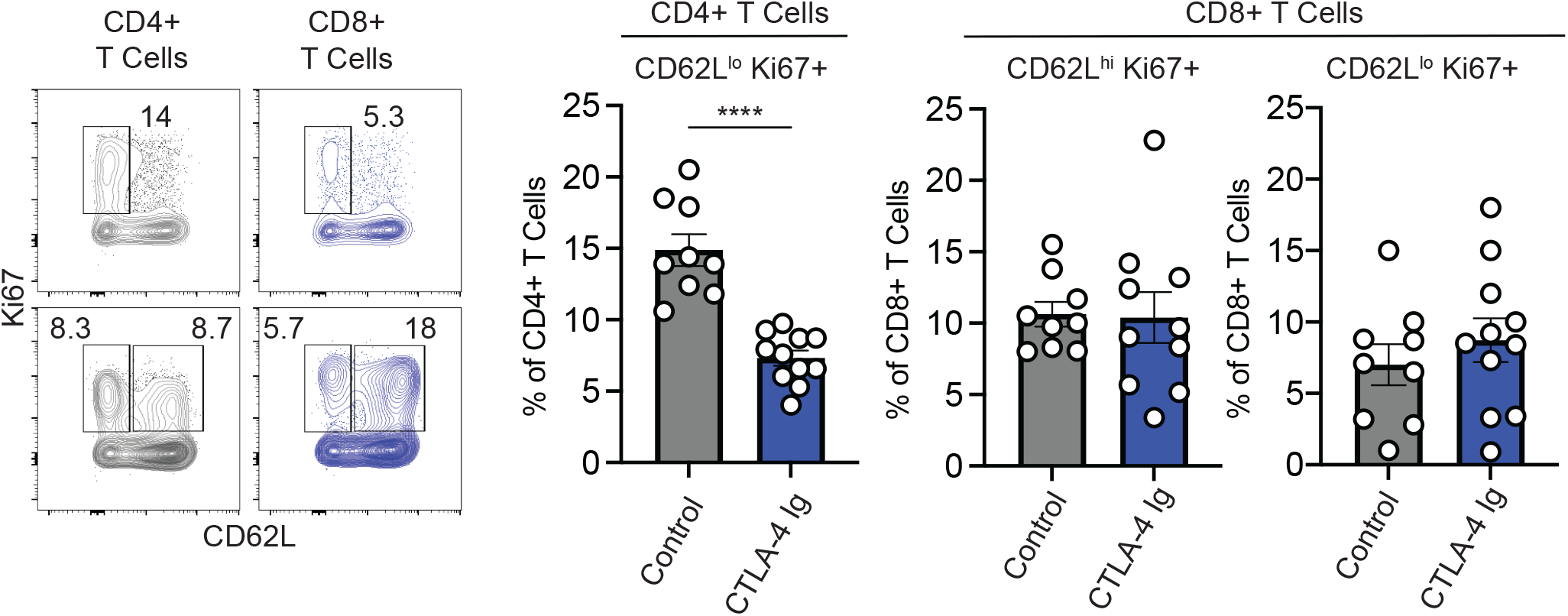
CTLA-4 Ig reduces the frequency of proliferating effector CD4^+^ T cells but not CD8^+^ T cells. Related to Figure 7. C57Bl/6 mice were grafted with H-2^d^ Balb/c skin and treated with CTLA-4 Ig or untreated. On day 14 after grafting, the phenotype of CD4^+^ and CD8^+^ T cells was assessed in the spleen. Each data point depicts an individual mouse. Summary data depicts pooled results from 2 independent experiments (9/group). Statistical analysis performed using unpaired Student’s t-test. Error bars depict SEM. ****p<0.0001.

